# Protein-peptide Interaction Representation Learning with Pretrained Language Models

**DOI:** 10.64898/2026.01.12.699173

**Authors:** Xinke Zhan, Silong Zhai, Tiantao Liu, Shaolong Lin, Tao Bi, Bingwen Zhu, Shirley W.I. Siu

## Abstract

Protein-peptide Interactions (PpIs) paly essential roles in diverse cellular processes, yet their systematic identification remains challenging due to the limited availability of experimentally annotated protein-peptide interaction data. To address this challenge, we present PepInter, a sequence-based Deep Learning (DL) framework that leverages large-scale pretraining on structurally derived pseudo protein-peptide pairs to learn interaction-relevant representations. Specifically, energy-dominant peptide fragments are extracted from protein complexes curated from non-redundant Protein Data Bank (PDB) structures, enabling the construction of pseudo protein-peptide interaction pairs that capture interface interaction patterns shared with canonical protein-protein interactions. This strategy allows the model to acquire interaction-aware priors in the absence of large-scale annotated protein-peptide complex datasets. Built upon the ESM-Cambrian (ESMC) architecture, PepInter adopts a two-stage pretraining strategy. In the first stage, masked language modeling is used to learn general protein sequence representations. In the second stage, the model is further trained to predict Rosetta-derived energetic scores, explicitly incorporating structural interaction signals into the learned embeddings. Following pretraining, PepInter is fine-tuned for both protein-peptide interaction classification and peptide bioactivity regression tasks. Across multiple benchmark datasets, including protein-peptide binding affinity prediction, PepInter consistently outperforms existing baseline methods and demonstrates strong generalization in identifying biologically meaningful PpIs. Case studies further highlight its ability to recover known interaction patterns and predict novel protein-peptide interactions. Together, these results establish PepInter as a scalable and effective framework for protein-peptide interaction prediction, with strong potential to accelerate peptide-based drug discovery.

## Introduction

The identification of potential interactions between peptides and proteins plays a crucial role in elucidating various biological functions, such as gene expression regulation and enzyme activity modulation [1]. Several studies have reported that the peptides exhibit high specificity and relatively low immunogenicity, making them attractive candidates for therapeutic applications [2–3]. Moreover, the biological activity of peptides is mainly achieved through their interactions with proteins [4]. Some peptides can bind to protein or cell surface receptors selectively for activating or inhibiting signal transduction [5]. Therefore, predicting peptide-protein interactions (PpIs) are essential for understanding cellular processes and guiding the design of peptide-based therapeutics. Traditional experimental methods have been widely used to detect PpIs. However, they are labor-intensive and time-consuming, requiring extensive efforts such as mutagenesis, binding assays and structural characterization, which limiting their large-scale application [6]. Predicting potential PpIs using computational approaches is an important and challenging research topic [7–8]. In recent years, *in silico* prediction has emerged as a powerful strategy to complement experimental efforts in characterizing protein-peptide interactions and predicting binding sites on protein surface [9–10]. The Computational strategies for predicting PpIs can be broadly categorized into two classes: molecular docking-based and machine learning-based approaches. Docking-based methods [11–13] perform global searches over protein surfaces to identify potential binding sites, followed by conformational sampling, docking, and selection of the optimal binding mode. Despite the methods are widely used in PpIs, there still exist several limitations, such as conformational sampling and handling of protein-peptide flexibility are insufficient, the performance of blind prediction scenarios with unknown binding sites prediction still needs to be improved.

Recently, the number of PpIs have been experimentally validated, which has driven the development of machine learning-based methods such as SPRINT-seq [14] and PepBind [15], while their dependence on handcrafted features and limited generalization capability hinders their applicability to more complex and diverse prediction scenarios. The rapid development of DL [16–17] has made large-scale prediction possible. For instance, Zhou et al. reported a multi-task framework named PepGPL [18], which fully consider the local interaction between peptides and proteins, it builds an interaction graph which shows strong ability in capturing the interaction information of global sequence and local residues. Sun et al. proposed a deep learning method based on convolutional neural network and multi-head attention called PepPAP, sequence-based features such as sequence encoding, interface propensity, physicochemical properties and intrinsic disorder shows great prediction performance [19]. With the rapid development of natural language processing (NLP) techniques, large language models (LLMs) such as ESM [20], BERT [21] and ProtTrans [22] have been introduced to extract high-dimensional embeddings from amino acid sequences. Jin et al. proposed a transformer-based model named TPepPro, which combine the features of local protein sequence and global structure, and optimized the BN-ReLU that notably reduced the computing resources, shows great potential in PpIs prediction [23]. Compared with docking-based methods, machine learning-based approaches are applicable to a broader range of proteins and peptides by learning informative sequence representations, thereby facilitating efficient interaction screening and the discovery of novel functional roles.

A major obstacle in developing DL methods for modeling PpIs is the limited available of high-quality training data. Experimentally validated PpIs datasets are typically small, sparsely distributed across protein families and biased toward a narrow set of well-studied targets, which poses several challenges for model training. Insufficient coverage of sequence and interaction space hinders the ability of models to learn generalizable interaction patterns, leading to overfitting and poor performance on unseen proteins or peptides. The lack of diverse interaction contexts limits the effectiveness of data-hungry architectures, particularly large sequence-based language models and graph-based neural networks, which rely on abundant and heterogeneous data to learn robust representations. As a result, models trained on limited experimental PpI data often fail to generalize beyond the training distribution and exhibit reduced predictive reliability. To mitigate these limitations, we draw inspiration from the “hot segment” paradigm of protein-protein interactions, the interaction between two proteins can be mediated by a linear segment from one protein that contributes most of the interface energy [24]. Based on this principle, we leverage protein-protein complex to systematically expand the training dataset by deriving pseudo protein-peptide interaction pairs, providing a natural and scalable data source for model pretraining. In addition, pretrained language models have been widely adopted to generate informative protein sequence representations, and their success has driven advances in sequence-based PpIs modeling.

In this study, we present PepInter, a sequence-based DL framework that integrates energy-guided structural priors with language model-based representation learning. Specifically, we collect protein-protein interaction data from the public databases and apply the Rosetta Peptiderive protocol [25] to extract energy-dominant peptide fragments, yielding large-scale pseudo protein-peptide pairs. These pairs are subsequently filtered to remove low-quality structures and fragments with negligible interface energy contributions, ensuring reliable interaction-relevant supervision. Energy-guided pretraining on structure-derived pseudo protein-peptide pairs endows PepInter with interaction-aware priors, which are further refined through ESMC-based masked language modeling (MLM) to integrate sequence semantics and latent interaction features. Subsequently, PepInter is fine-tuned on two complementary experimentally annotated datasets. The protein-peptide interaction dataset provides supervision for binary PpI classification, whereas the protein-peptide binding affinity dataset further refines the learned representations by modeling quantitative interaction strength, thereby ensuring that PepInter captures both interaction specificity and binding potency. Extensive evaluations using five-fold cross-validation and independent test sets demonstrate that PepInter consistently outperforms state-of-the-art methods on benchmark datasets, highlighting its effectiveness as a powerful tool for large-scale PpI prediction.

## Materials and Methods

### Overview of PepInter

Protein-peptide interactions are governed by sequence patterns that encode both biochemical compatibility and implicit structural constraints. However, learning such interaction-relevant sequence representations is challenging due to the limited availability of experimentally annotated protein-peptide data. To address this challenge, we propose PepInter to learn transferable protein-peptide interaction representations that can support diverse downstream tasks, such as interaction identification and binding affinity prediction. To achieve this goal, PepInter leverages large-scale structure-derived pseudo protein-peptide data to inject interaction-aware priors into sequence representation learning and subsequently adapts these representations to specific prediction objectives using limited experimental annotations. The overall architecture of PepInter is illustrated in Figure 1. The framework consists of four parts: data collection; data generation; model pretraining and fine-tunning; downstream task. PepInter adopts a residue-level MLM strategy based on the ESMC architecture and is pretrained on the constructed pseudo protein-peptide pairs. This pretraining procedure encourages the model to encode sequence patterns that are statistically associated with protein-protein interface regions and energy-dominant peptide segments, thereby transferring structural interaction knowledge into the learned sequence representations. For downstream tasks, PepInter is adapted to different prediction objectives by attaching lightweight task-specific output modules and fine-tuning the pretrained backbone on experimentally validated protein-peptide datasets. These tasks include discriminating interacting from non-interacting protein-peptide pairs as well as estimating quantitative binding strength, enabling PepInter to address both interaction specificity and interaction potency within a unified modeling framework. By combining large-scale pseudo protein-peptide pretraining with effective transfer of interaction-relevant structural priors, PepInter substantially improves predictive performance on real-world protein-peptide interaction identification and binding affinity estimation tasks.

**Fig. 1.**
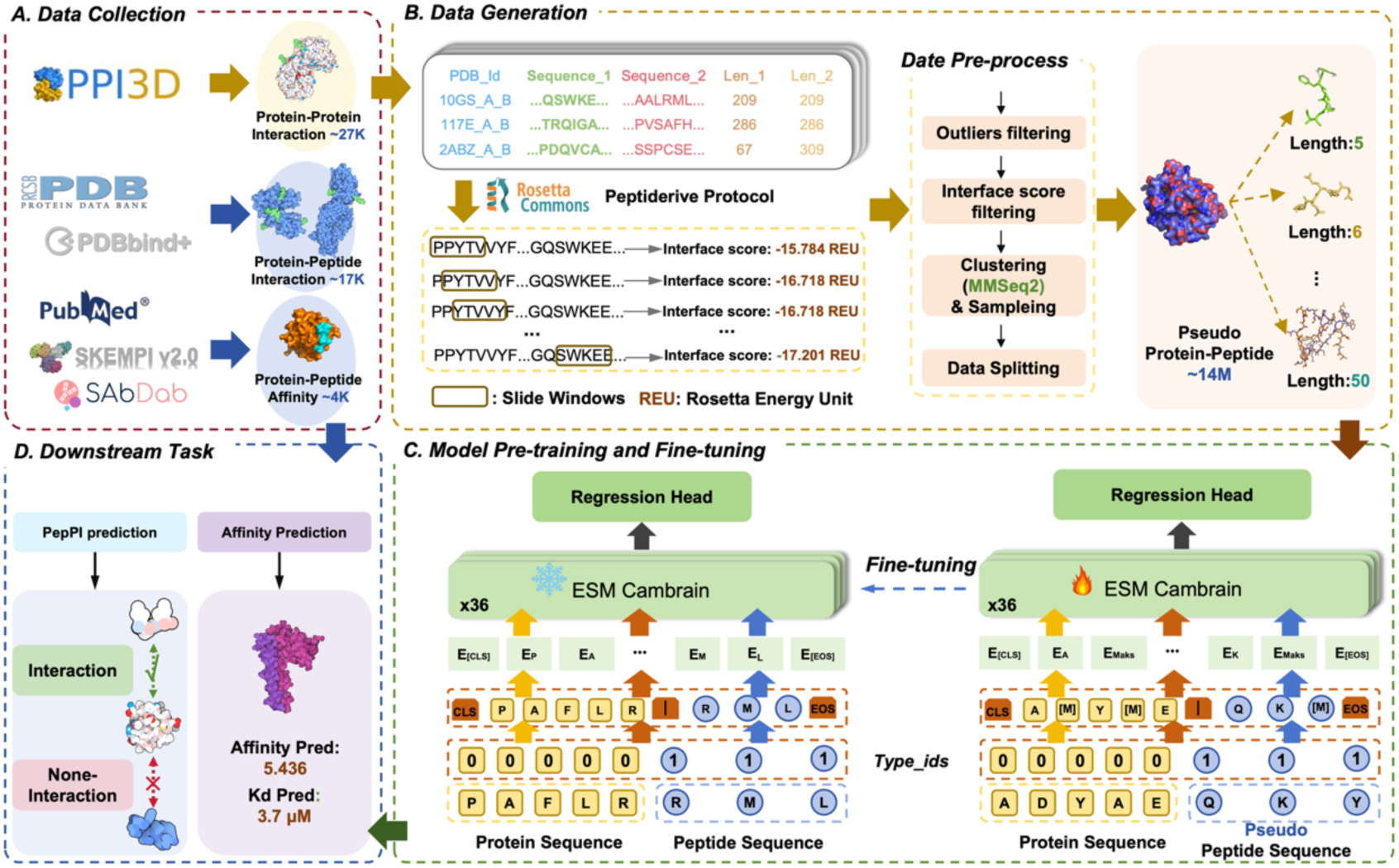
The workflow of PepInter. A) **Data collection:** protein-protein interaction data, protein-peptide affinity data, and protein-peptide interaction data are collected from several databases; B) **Data generation:** large-scale pseudo peptide-protein pairs are generated from protein complexes using the Rosetta Peptiderive protocol for subsequent pretraining; C) **Model pre-training and fine-tuning:** the ESM-C model is adopted as the encoder backbone, and residue-level MLM is performed on the pseudo data pairs to learn implicit structural features within peptide-protein interaction space. For downstream tasks, PepPI and Affinity prediction heads are added and fine-tuned using experimentally validated peptide-protein data to adapt to specific prediction objectives; D) **Downstream task:** PepInter enables both peptide-protein interaction prediction and binding affinity estimation.

### Data curation

#### Pseudo protein-peptide pretraining dataset-ProtFragDB

ProtFragDB is constructed using protein complexes from the PPI3D database [26] which contains more than 891,000 experimentally determined PDB Protein-Protein Interaction (PPI) pairs. Several filtering criteria were applied: (i) a structural resolution less than 4.0 Å; (ii) a protein sequence length of fewer than 1000 amino acids; and (iii) a sequence identity threshold of 40% to reduce redundancy among PPI pairs. MMseqs2 [27] was employed to cluster all protein sequences using a 40% sequence identity threshold. After filtering and clustering, a total of 28,916 non-redundant PPI pairs. Subsequently, the Rosetta Peptiderive protocol was applied to these PPIs to generate pseudo protein-peptide fragment pairs. For each protein-protein complex, one chain was fixed as the receptor protein, while the interacting partner chain was systematically decomposed into linear peptide fragments using a sliding-window strategy with fragment lengths ranging from 5 to 50 amino acids [28]. For each generated pseudo peptide-protein pair, Peptiderive computed an interface energy score to quantify the energetic contribution of the peptide fragment to the interaction interface. The pairs associated with abnormal or erroneous interface scores were discarded. To further eliminate redundancy, the resulting pseudo peptide-protein pairs were clustered again using MMseqs2 with a 40% sequence identity threshold. Sequences belonging to the same cluster were strictly assigned to the same data split to prevent information leakage between the training and validation sets. After deduplication, approximately 9 million high-quality pseudo peptide-protein fragment pairs were retained, including 8,983,271 pairs for training and 22,117 pairs for validation.

#### Protein-peptide interaction dataset-PepInterDB

The protein-peptide interaction dataset namely PepInterDB was constructed by systematically scanning all protein complex structures deposited in the RCSB Protein Data Bank (PDB, accessed in September 2025) [29]. A Python-based pipeline was developed to identify interacting chain pairs, where two chains were considered to form an interaction if any pair of heavy atoms between them was within a distance threshold of 5 Å. To specifically capture protein-peptide interactions, interacting chain pairs were further filtered by chain length, requiring the protein chain to contain 40-1000 residues and the peptide chain to contain 5-40 residues. Redundant interaction pairs with identical protein-peptide sequence combinations were removed. Following this procedure, a total of 8,527 non-redundant protein-peptide interaction pairs were obtained. The dataset was randomly split into a training set containing 6,630 samples and a validation set containing 1,897 samples. An equal number of negative samples were additionally generated, yielding a final dataset of 17,054 protein-peptide pairs. The protein and peptide length distributions of PPInterDB are shown in Figure 2(A) and Figure 2(B), respectively. Figure 2(C) compares the amino acid composition of protein and peptide sequences, revealing broadly similar distributions with subtle differences across residues. In both above datasets, sequences containing non-standard amino acids were retained provided that the number of such residues did not exceed half of the total sequence length; otherwise, the corresponding samples were removed. This strategy was adopted to maximize data utilization while maintaining overall sequence quality. Notably, non-standard amino acids were allowed only in the training set, whereas the validation set contained exclusively standard amino acid residues. This design intentionally introduces mild noise during training, which is expected to improve model robustness and generalization performance.

**Fig. 2.**
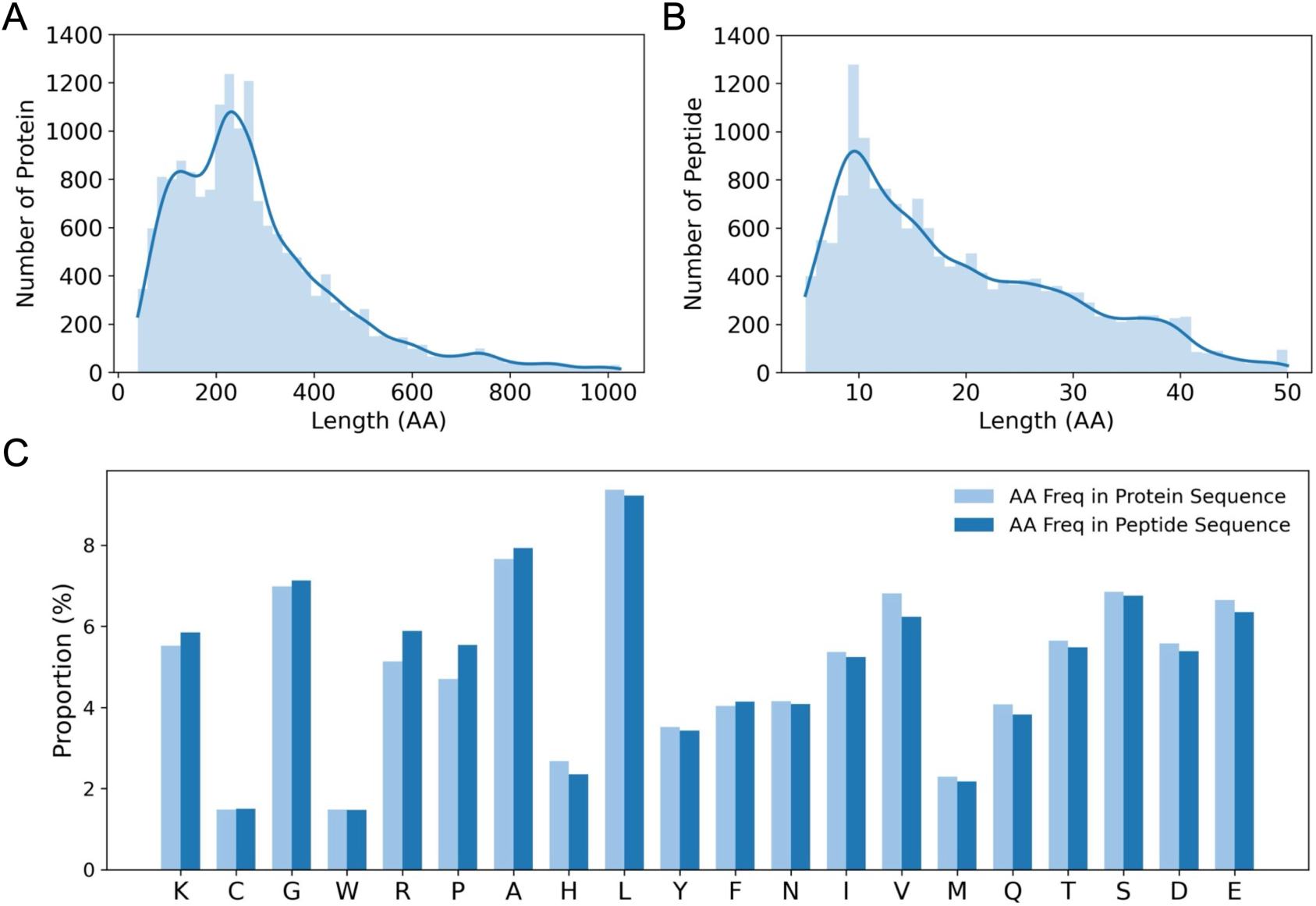
Overview of the characteristics of PepInterDB (A) Length distribution of protein sequences in PepInterDB. (B) Length distribution of peptide sequences in PepInterDB. (C) Comparison of amino acid composition between protein and peptide sequences in PepInterDB.

#### Protein-peptide affinity dataset-PepAffDB

Several works have previously reported protein-peptide affinity dataset, but the existing datasets are often incomplete in both quality and scale. To better evaluate the generalization ability of the proposed model, a protein-peptide affinity dataset PepAffDB was constructed that collected from multiple sources, including PDBbind v2020R1 [29], SKEMPI v2.0 [30], SabDab [31], UmamiMeta [32] and published literature. Specifically, a total of 937, 246, 2,069 and 1,869 protein-peptide affinity records were collected from PDBbind v2020R1, SabDab, UmamiMeta and SKEMPI v2.0, respectively. In addition, 481 protein-peptide records were manually curated from more than 500 published articles through a systematic literature search using keywords related to “protein-peptide affinity”. These records include experimentally synthesized peptides, sequence variants and other peptide modifications, most of them do not have resolved structural information. After integrating all sources, extensive data cleaning, filtering and deduplication were performed, resulting in 4,077 affinity records. To ensure data reliability, peptides shorter than 5 residues were excluded, yielding a final dataset of 3,670 high-quality affinity pairs. Figure 3(A) summarizes the changes in the number of protein-peptide affinity pairs and Figure 3(B) presents the distribution of binding affinities in the final PepAffDB dataset. Moreover, the PepAffDB were clustered using MMseqs2 with an 80% sequence identity threshold. Sequences belonging to the same cluster were strictly assigned to the same data split to prevent information leakage, resulting in 2,698 pairs for training, 671 pairs for validation, and 301 pairs for testing. Affinity measurements reported in different units, including *K_d_*, *K_i_*, and *IC*_50_, were retained as provided by the original sources. The affinity values were converted into a unified logarithmic scale to facilitate model training and evaluation, while records lacking sufficient information for reliable conversion were excluded. The final dataset covers a broad range of binding strengths spanning multiple orders of magnitude, enabling robust assessment of model performance across diverse affinity regimes. The dataset can be obtained in github.

**Fig. 3.**
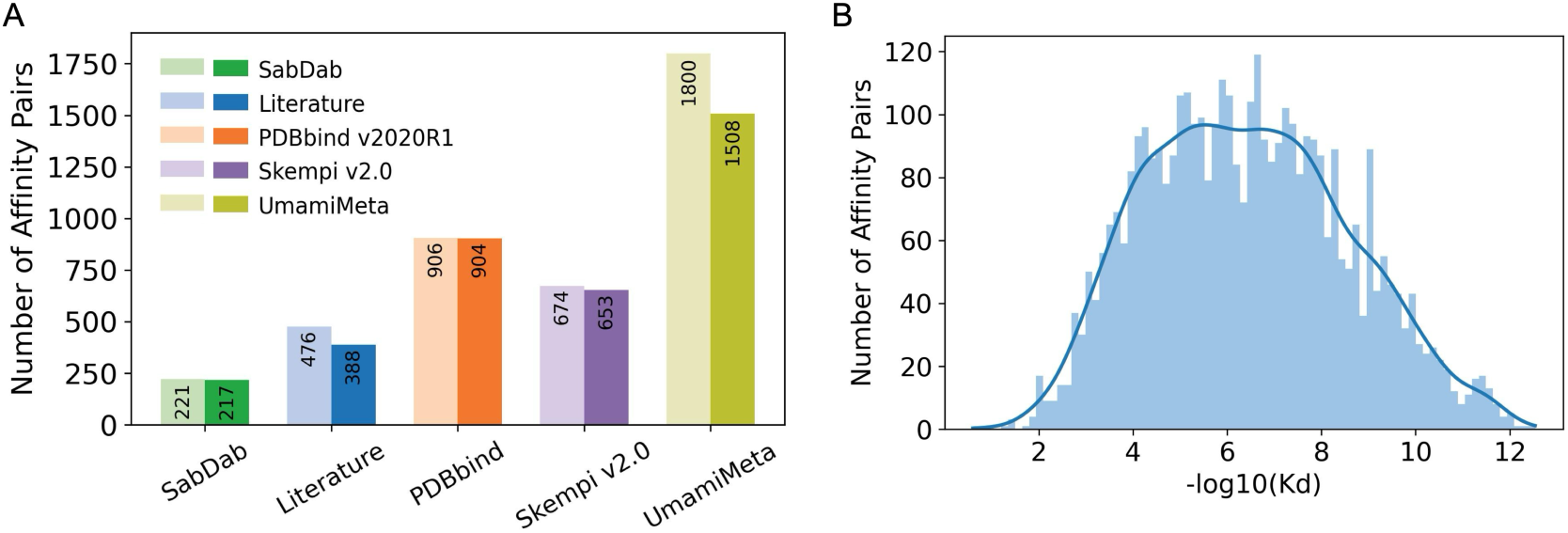
Overview of the characteristics of PepAffDB. (A) Changes in the number of protein-peptide affinity records in PepAffDB after excluding peptides shorter than 5 residues across different data sources. (B) Distribution of binding affinities in the final curated PepAffDB shown in − log*_di_*(*K_d_*) space.

#### Cluster-based data splitting strategy

Due to a peptide can likewise bind to multiple similar proteins and a protein can also interact with multiple similar peptides, this pattern often contains substantial redundancy. To eliminate the potential bias introduced by such redundancy and to ensure a fair comparison, cluster-based validation strategy is proposed [33]. Specifically, based on the protein and peptide sequences in the dataset, we employed MMSeqs2 to perform clustering of proteins, peptides, and protein-peptide pairs under different sequence similarity thresholds (0.3, 0.5, and 0.8), and constructed three evaluation settings, namely “Novel Protein”, “Novel Peptide”, and “Novel Pair”. In the “Novel Protein” setting, clustering was performed on protein sequences, whereas in the “Novel Peptide” setting, peptide sequences were clustered independently. The “Novel Pair” setting involved clustering protein-peptide pairs. Based on the clustering results, the dataset was divided into training and test sets at an 8:2 ratio, with the primary constraint that samples belonging to the same cluster were not allowed to appear simultaneously in both the training and test sets, thereby ensuring the validity and fairness of the experimental evaluation.

### Model architecture

#### Model Pre-training

The quality of protein representations directly determines the performance of downstream prediction tasks. Recently, protein large language models (LLMs) trained in an unsupervised manner on massive sequence datasets such ESM series have become base tools for obtaining high-quality protein embeddings. The ESM-C [34] has demonstrated remarkable advantages in representation learning that it outperforms ESM-2 while offering faster inference speed and lower computational cost with a comparable number of parameters. The 600 M-parameter ESM-C achieves computational efficiency comparable to that of the 300 M-parameter ESM-2 model. Given this advantage, ESM-C is adopted as the base architecture during the pre-training stage, with a masked language modeling (MLM) strategy employed as the self-supervised learning objective on protein and peptide sequences. Given a group of protein-peptide pair:

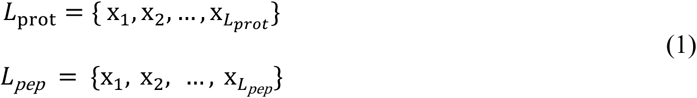

where *L_prot_* means the length of protein sequence and *L_pep_* denotes the length of peptide sequence. We define a template processing of the tokenizer to standardize the format of input sequences, enabling the model to clearly distinguish protein sequences and peptide sequences. Specifically, we defined a protein-peptide pair template as follows:

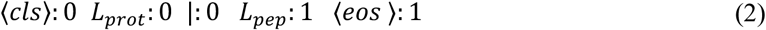

where 0 and 1 mean the token-type ID for distinguish the protein and peptide sequence, respectively. The symbol ‘|’ is defined as a delimiter connecting the two sequences. All special symbols in the template (<CLS>, <EOS>, ‘|’) are registered in the vocabulary as special tokens and are converted into their corresponding token IDs. This design primarily serves the following purposes: The start and end markers are added to the input, enabling the model to explicitly identify sequence boundaries; The token-type IDs differentiate the protein and peptide components, and the delimiter clearly marks their separation. These operations help the model better understand the definition of a protein-peptide pair and improve input consistency and stability in downstream prediction tasks. During training, approximately 15% of amino acid residues are randomly selected as masked positions, and the model is trained to predict the true amino acid at each masked site, learn semantic and structural patterns within the sequences. The training loss is defined as follows:

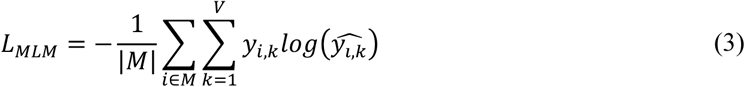

where |*M*| denotes the number of masked tokens; *V* means the size of the vocabulary; *y_i,k_* is an indicator function that equals 1 if the true class of token *i* is *k*, and 0 otherwise; 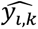 is the probability predicted by the model that token *i* belongs to class *k*. The model can automatically capture amino acid co-occurrence preferences, local and long-range dependency patterns, as well as structural and functional information implicit in protein-peptide interactions via a self-supervised objective, thereby providing more generalizable sequence representations for downstream prediction tasks.

#### Fine Tunning

During the fine-tuning stage, the model is initialized with the parameter weights learned during pretraining which fully leveraging the general representation capabilities acquired from large-scale protein-peptide sequence pairs. The model is then trained in a supervised manner on the protein-peptide dataset. The inputs are the protein sequence *L_prot_* and the peptide sequence *L_pep_*. Depending on the task setting, the prediction *ŷ* represents either a binary classification probability, or a continuous binding affinity value. For the classification task, the Focal loss [35] is adopted which is a modified cross-entropy loss function tackles extreme class imbalance, the original loss function is shown as follows:

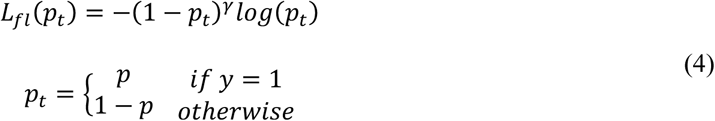

where *γ* is set to 2. In our experiment, the *⍺*-balanced variant of the focal loss is employed:

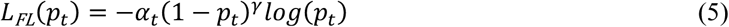

where *⍺*_1_ is set to 0.25.

For the binding affinity prediction task, a Mixture Density Network (MDN) [36] loss is adopted to model the potentially multi-modal distribution of binding affinities. Specifically, the MDN head outputs the parameters of a *K*-component Gaussian mixture, including mixture weights *π* ∈ *R^K^*, means *μ* ∈ *R^K^*^×*D*^, and standard deviations *σ* ∈ *R*^K×*D*^. The training objective is the negative log-likelihood:

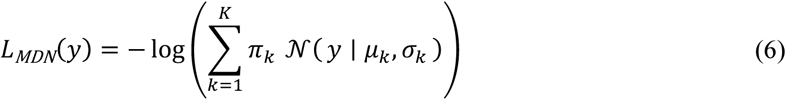

To improve robustness against label noise and hard outliers, we further apply a small-loss selection strategy during training. For each mini-batch, we first compute the per-sample MDN loss 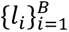 , and determine a quantile threshold *q_r_* at a predefined retention ratio *r* (e.g., r = 0.8). Only samples with *l_i_* ≤ *q_r_* are assigned non-zero weights and used to update the model, with the weights normalized to keep the average weight close to 1. The final training objective is:

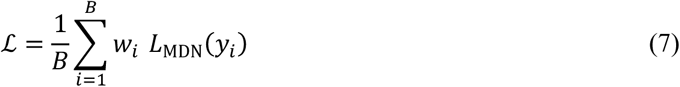

where w_F_ ∈ {0,1} indicates whether the *i-th* sample is selected. This small-loss trick adaptively filters out high-loss samples which are potentially noisy, stabilizing training and yielding more robust estimates of the protein-peptide behavior.

#### Prediction Head of PpI and Affinity

The protein-peptide interaction prediction task aims to determine whether a given protein-peptide pair exhibits an interaction relationship, mapping the input sequence pair to a binary label {0, 1}. In this task, The classification head is implemented as a two-layer feed-forward network. Given a feature representation 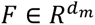, the hidden representation is computed as:

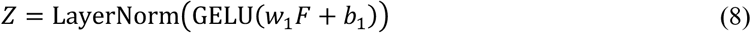

where GELU is the activation function, which provides a smooth, non-linear transformation. The final output logits are then obtained by:

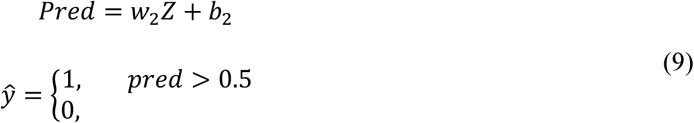

For binary interaction prediction, a sigmoid function is applied to produce the interaction probability *ŷ*.

For the binding affinity prediction task, we employ a MDN head to model the potential binding affinities. Given a feature representation 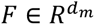, the final training loss is computed as a normalized weighted average over the selected samples. At inference time, the predicted binding affinity is obtained as the expectation of the mixture distribution:

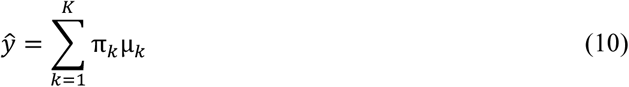

where *K* denotes the number of Gaussian components, *π*_5_ represents the mixture weight of the *k-th* component, and *μ*_5_ denotes the corresponding component mean. The predicted binding affinity *ŷ* is computed as the expectation of the mixture distribution.

### Evaluation and Configuration

#### Evolution Metrics

This study employs widely used binary classification metrics to comprehensively evaluate the performance of various machine learning models in PpIs classification tasks. These metrics include accuracy (ACC), recall (REC), F1-score, the area under the ROC curve (AUC), and the area under the precision-recall curve (AUPR). Overall, these metrics not only assess the overall predictive performance but also evaluate their ability to maintain balance between positive and negative classes. The definitions of these metrics are as follows:

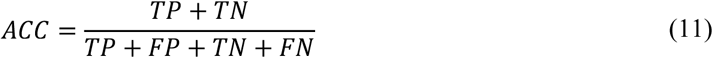

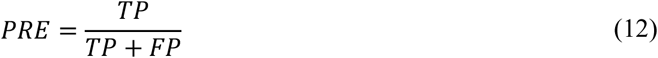

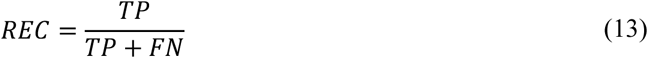

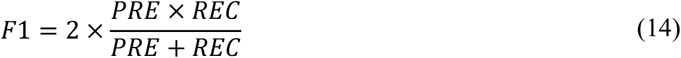

In this context, True Positive (TP) and True Negative (TN) represent the number of positive samples correctly identified by the model, the number of negative samples that are correctly classified, respectively. False Positive (FP) and False Negative (FN) refer to negative samples that are incorrectly predicted as positive, positive samples that are incorrectly classified as negative, respectively.

Similarly, the root mean square error (RMSE), mean absolute error (MAE), and coefficient of determination (R²) are used in the protein-peptide affinity regression task. RMSE and MAE reflect the absolute magnitude of prediction errors, with smaller values indicating better performance; R² measures the explanatory power of the model, and values closer to 1 indicate better fitting capability. Their definitions are as follows:

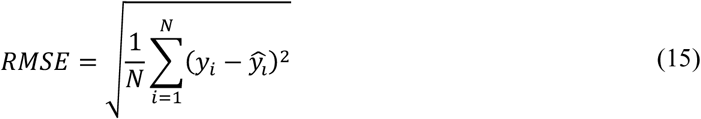

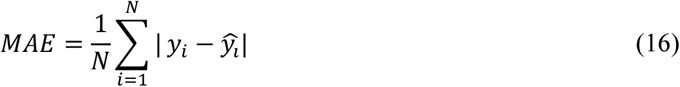

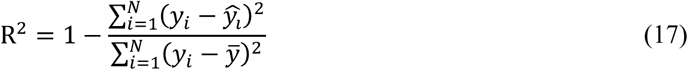

where *y_i_* and *ŷ_i_* denote the true and predicted values, respectively; *ȳ* represents the mean of the true values, and *N* is the number of samples.

#### Hyperparameter and Training Configuration

In the construction of ProtFragDB, the Rosetta Peptiderive protocol was employed to identify protein-protein interaction interfaces from given protein structures and to predict peptide fragments with potential interface-binding capability. In Peptiderive, the primary parameters “*restrict_receptors_to_chains*” and “*restrict_partners_to_chains*” are specified to define the receptor protein chains and the corresponding partner chains from which peptides were derived. The option “*dump_peptide_pose*” is set to true to output the conformations of the derived peptide fragments, while “*dump_prepared_pose*” is enabled to retain the preprocessed protein-peptide complex structures prior to docking. All remaining parameters were kept at their default values.

During the pre-training phase, 36 Transformer layers are employed with each layer containing 18 attention heads, the model is trained for three steps with a batch size of 128. the maximum sequence length is set to 1000 for proteins and 50 for peptides. The hidden dimension is 1152 and AdamW [37] is used as the optimizer with an initial learning rate of 1 × 10^HI^. The maximum number of training steps is 10,000. Pre-training is conducted on three NVIDIA A800 80GB PCIe GPUs and converges in approximately 72 hours.

The model is fine-tuned on a real protein-peptide interaction dataset PepInterDB. Protein and peptide sequences are encoded separately using the pre-trained ESM-C encoder, the resulting residue-level contextual representations are subsequently fused and fed into a binary classification prediction head to output the binding probability. Full-parameter fine-tuning is adopted so that the pre-trained representations can be effectively transferred to the downstream task. The batch size is set to 24, the maximum number of prediction epochs is 5, and the learning rate is set to 1 × 10^HI^. Similarly, in the protein-peptide affinity prediction stage, we fine-tune the model on a real protein-peptide affinity dataset PepAffDB. The fused representations are fed into a regression prediction head to output the binding affinity. The fine-tuning configuration is consistent with that used in the interaction prediction stage, with a maximum of 50 prediction epochs and a learning rate of 1 × 10^HI^. The detailed of comparison methods are listed in supplementary file.

## Results

### Comparative performance of PepInter against the baseline methods

To better evaluate the performance of our proposed method, PepInter is compared with several competitive approaches. All baseline methods were locally installed, trained and tested on the same datasets use five-fold cross-validation to ensure fairness. The baseline methods include XGBoost [38], MAARDTI [39], HyperAttentionDTI [40], and Rep-ConvDTI [41]. We trained multiple baseline models on our self-constructed dataset, the prediction results are presented in Table 1, where the best-performing method for each metric is highlighted in bold.

**Table 1.** Performance of PepInter against five methods using five-fold cross-validation on our own dataset.

| Models | ACC | REC | F1 | AUC | AUPR |
| --- | --- | --- | --- | --- | --- |
| XGBoost | <u>0.803±0.002</u> | <u>0.636±0.005</u> | <u>0.764±0.003</u> | <u>0.931±0.003</u> | <u>0.938±0.003</u> |
| MAARDTI | 0.757±0.014 | 0.548±0.035 | 0.692±0.260 | 0.873±0.005 | 0.893±0.005 |
| PIPR | 0.716±0.011 | 0.520±0.021 | 0.646±0.018 | 0.806±0.014 | 0.816±0.013 |
| Rep-ConvDTI | 0.763±0.006 | 0.570±0.016 | 0.706±0.011 | 0.853±0.006 | 0.877±0.004 |
| HyperAttentionDTI | 0.765±0.014 | 0.568±0.038 | 0.755±0.017 | 0.883±0.009 | 0.899±0.006 |
| **PepInter** | **0.853±0.006** | **0.747±0.005** | **0.836±0.009** | **0.940±0.005** | **0.947±0.004** |

PepInter achieves the best performance across all evaluation metrics in the five-fold cross-validation experiments. The PepInter delivers stable improvements compared with the second-best model in each category. We can observe that our model achieves an improvement of 5%, 11.1%, 7.2%, 9%, 0.9%, 0.9% in ACC, REC, F1, AUC, AUPR, respectively, over the baseline model, XGBoost. The improvements reflect our model exhibits stronger robustness in distinguishing positive and negative samples and in identifying rare interaction pairs. PepInter achieves a higher recall and therefore a better overall performance, indicating that the model is more effective in maximizing the identification of true interaction pairs. These results indicate that PepInter is capable of capturing semantic relationship between protein and peptide sequences, substantially enhancing protein-peptide interaction prediction accuracy.

### Comparative performance of PepInter against baseline methods under different clustering settings

To assess the effectiveness of the proposed method, we compared PepInter against five baseline methods. The evaluation was conducted on three benchmark data set scenarios: “Novel protein”, “Novel peptide” and “Novel pair”. Model performance was quantified using ACC, F1 and AUC, respectively. The binary classification performance of all models is summarized in Figure 4. The PepInter consistently outperformed the baseline methods across all thresholds (NA, 0.3, 0.5, 0.8) in all three scenarios. We calculated the average performance of each method across four thresholds in each setting. Across all three evaluation scenarios, PepInter consistently outperforms the strongest baseline methods, including XGBoost and other deep learning-based approaches. In the most challenging “Novel Pair” setting, where both proteins and peptides in the test set are unseen during training, PepInter demonstrates the most pronounced advantage. Compared with the best-performing baseline, PepInter achieves up to 4.9% improvement in recall and 4.1% improvement in F1 score, indicating a substantially enhanced ability to identify true protein-peptide interactions under strict novelty constraints. In the “Novel Protein” setting, PepInter also exhibits clear superiority, particularly in recall, with improvements ranging from approximately 4.7% to 11.7% over the strongest baseline across different clustering thresholds. This suggests that PepInter is more effective at generalizing to unseen protein sequences while maintaining balanced overall performance. In the relatively less challenging “Novel Peptide” setting, PepInter continues to achieve consistent gains over baseline models, with improvements of approximately 2.0-2.9% in F1 score and 1.2-2.5% in recall, demonstrating robust and stable performance even when peptide sequences are novel. As a result, these results demonstrate the robustness of PepInter under varying degrees of sequence novelty and underscore its superiority over existing methods in binary protein-peptide interaction prediction tasks. The detailed results are provided in the Supplementary Information.

**Fig. 4.**
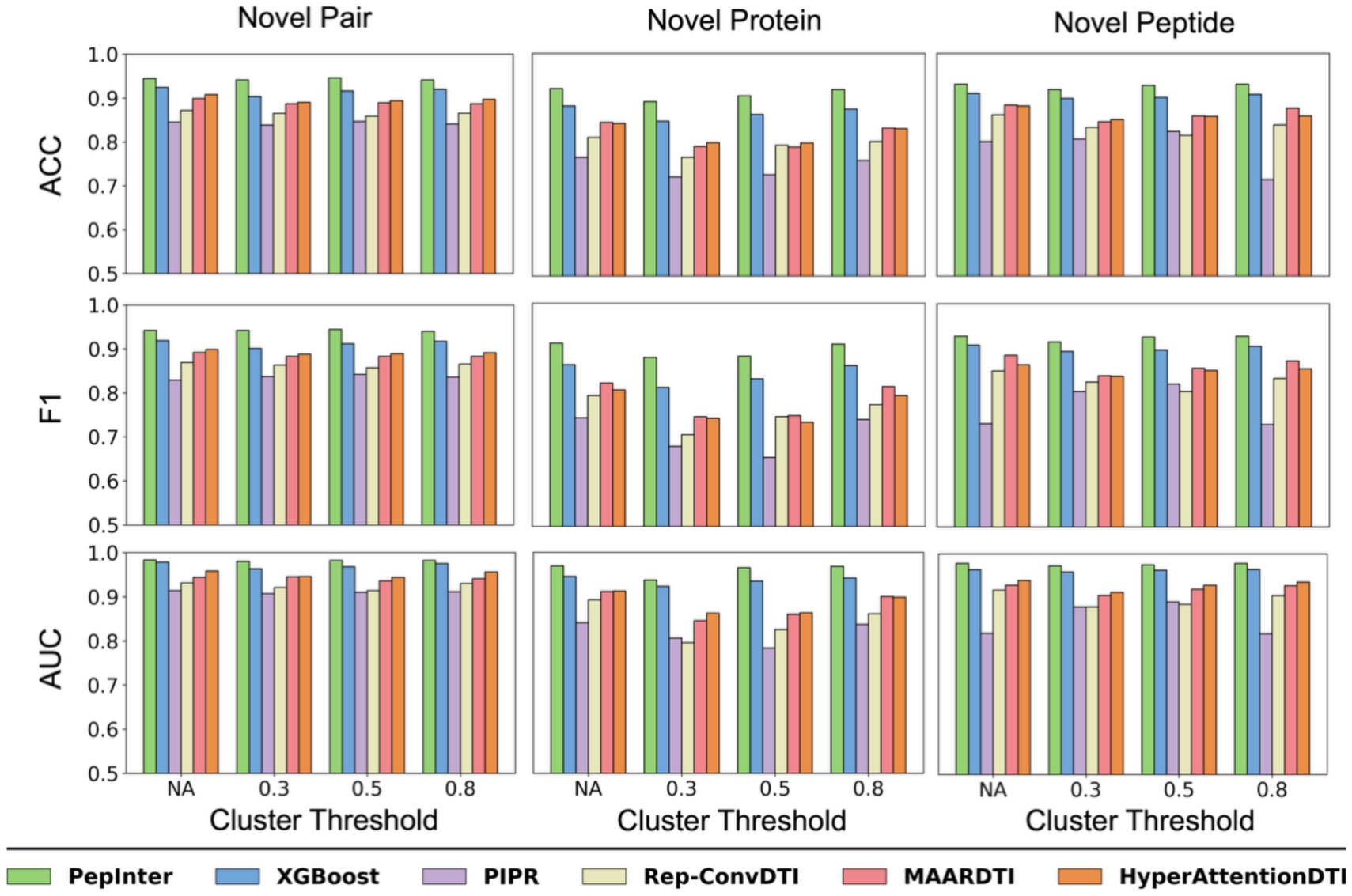
Performance comparison of PepInter and baseline methods under different clustering-based evaluation scenarios. The models are evaluated under “Novel Pair”, “Novel Protein”, and “Novel Peptide” settings across different sequence similarity thresholds (NA, 0.3, 0.5, and 0.8). “NA” indicates data sets not divided based on sequence similarity; Performance metrics include the ACC, F1 and AUC.

### Evaluation of Generalization Ability on Imbalanced Datasets

To evaluate the robustness of PepInter in binary interaction prediction and to simulate the scarcity of positive samples commonly observed in real biological experiments, we constructed imbalanced data distributions in the training set while keeping the original distribution unchanged in the test set. This ensures that the evaluation results faithfully reflect the model’s performance in realistic application scenarios. Under different positive-to-negative sample ratios, we compared the prediction performance of PepInter with several baseline models on these additional training sets. Specifically, we randomly paired peptides and proteins in the training set to generate negative samples for which no known interaction evidence exists. Four imbalanced datasets with ratios of 1:4, 1:9, 1:19, and 1:49 were generated. All models were trained on the full benchmark dataset.

As shown in Figure 5, PepInter consistently achieves the best performance across all test scenarios, demonstrating superior robustness compared with baseline models. As the positive-to-negative ratio decreases from 1:2 to 1:50, all models exhibit a downward trend in accuracy, indicating that severe class imbalance increases the difficulty of learning, particularly for baseline methods (Figure 5A). Nevertheless, PepInter maintains the highest accuracy and AUPR across all imbalance settings. As shown in Figure 5B, most models exhibit a slight improvement in AUPR when the ratio changes from 1:2 to 1:10, likely because additional negative samples provide richer discriminative information. However, under extreme imbalance conditions, models such as XGBoost and Rep-ConvDTI suffer from substantial performance degradation. In contrast, PepInter demonstrates consistently strong performance even when positive samples are highly sparse, highlighting its superior robustness and generalization ability. The strong performance of PepInter under severe class imbalance can be attributed to its ability to learn context-aware and interaction-focused representations. PepInter explicitly models residue-level interactions between protein and peptide sequences, enabling fine-grained characterization of potential binding relationships. As illustrated in Figure 6, the residue-level attention maps learned by PepInter exhibit clear and structured patterns, with interaction signals concentrated along specific regions rather than being diffusely distributed across the entire protein-peptide interaction space. This indicates that the model selectively emphasizes a small number of informative residues pairs and suppresses background noise arising from non-interacting regions. Such focused interaction patterns imply that the model learns meaningful contextual relationships between protein and peptide sequences, instead of relying on global sequence similarity alone. In the context of severely imbalanced data, where negative samples overwhelmingly outnumber positive ones, this interaction-focused representation is particularly advantageous. By concentrating predictive capacity on a restricted set of interaction-relevant residues, PepInter effectively reduces the influence of abundant non-informative negative samples. Consequently, the model is able to preserve discriminative binding signals even when positive interaction examples are extremely limited.

**Fig. 5.**
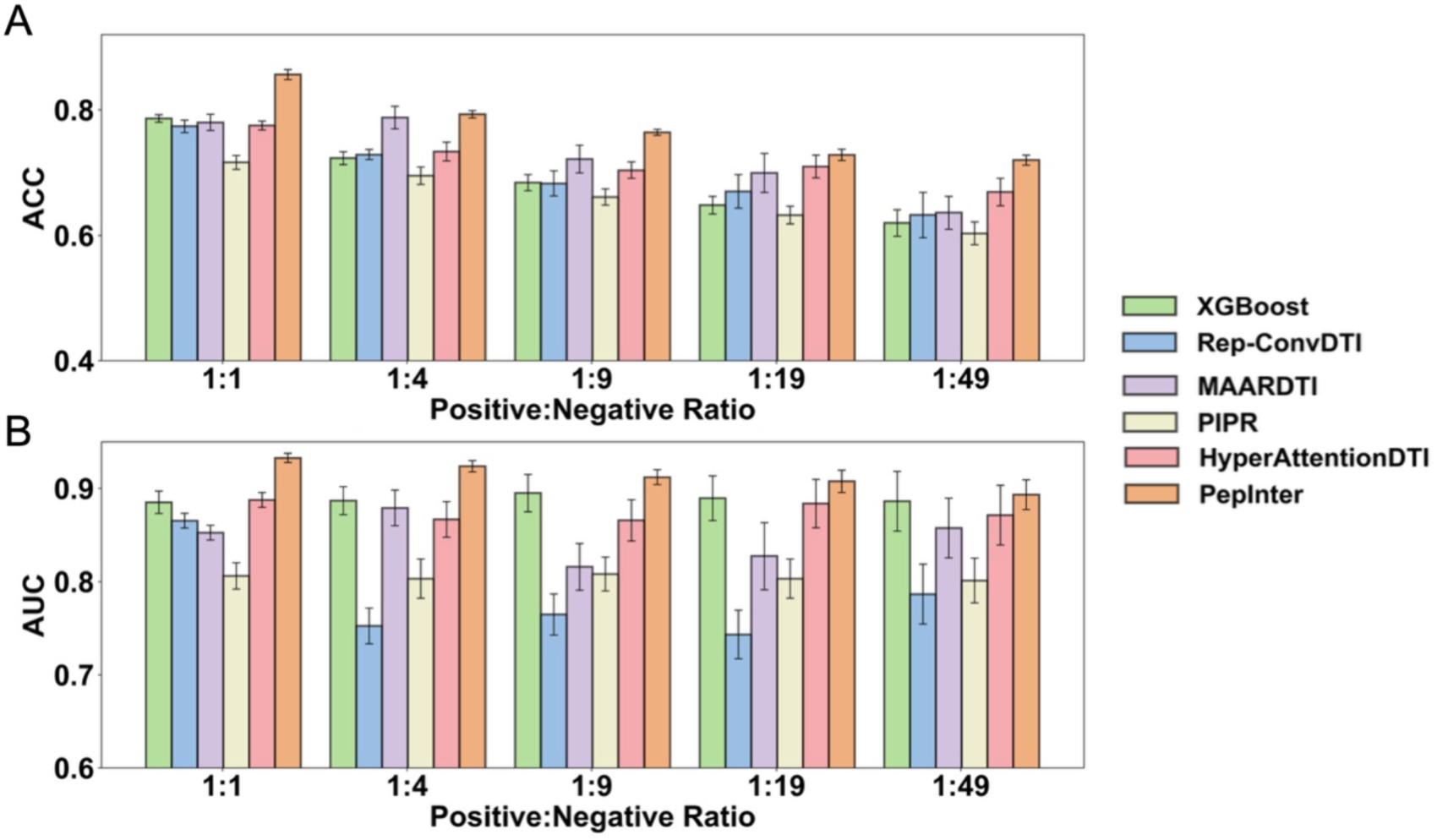
The PepInter outperformed the baseline models on an imbalance set. **A, B** show the evaluation results with different positive-negative ratios of the data set in terms of Accuracy and AUPR, respectively.

**Fig. 6.**
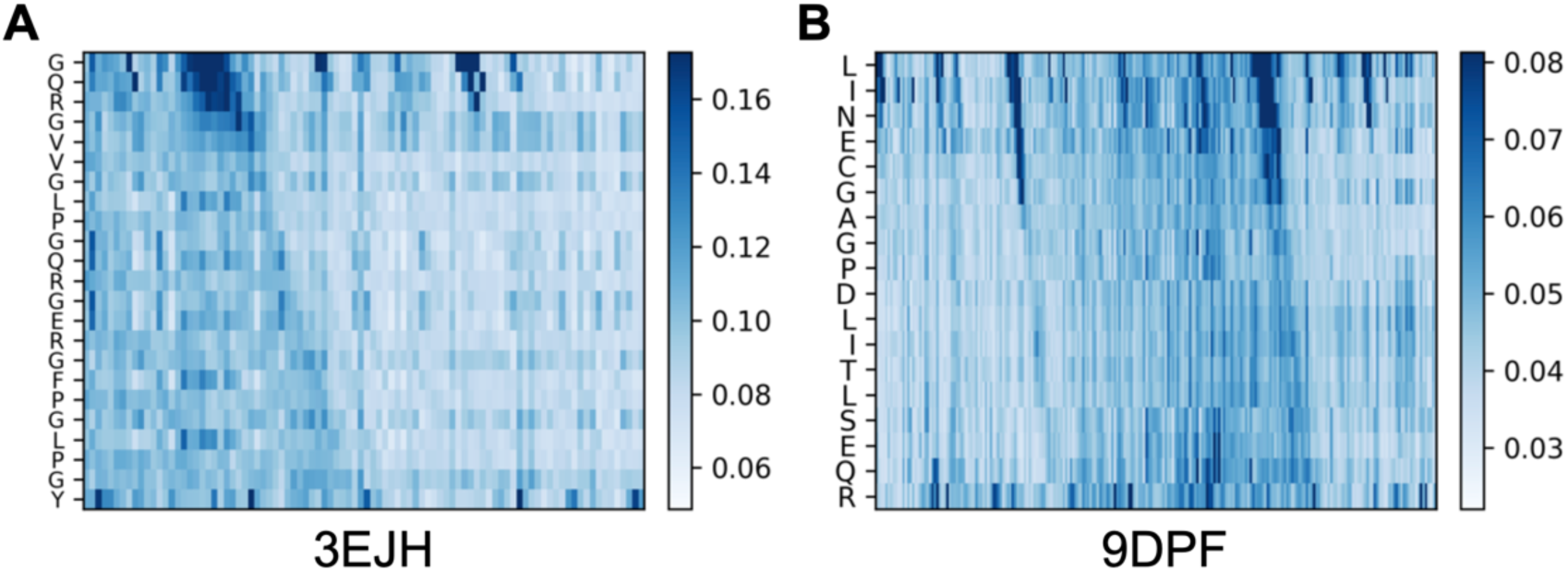
Visualization of interaction-focused representations learned by PepInter. (A) and (B) is attention map for protein-peptide complex derived from PDB: 3EJH and PDB: 9DPF. In each panel, the interaction signals are concentrated within specific protein and peptide regions rather than being uniformly distributed, indicating that PepInter captures localized and structured residue-level interaction patterns. The highlighted regions correspond to protein (orange) and peptide (green) segments, respectively.

**Fig. 7.**
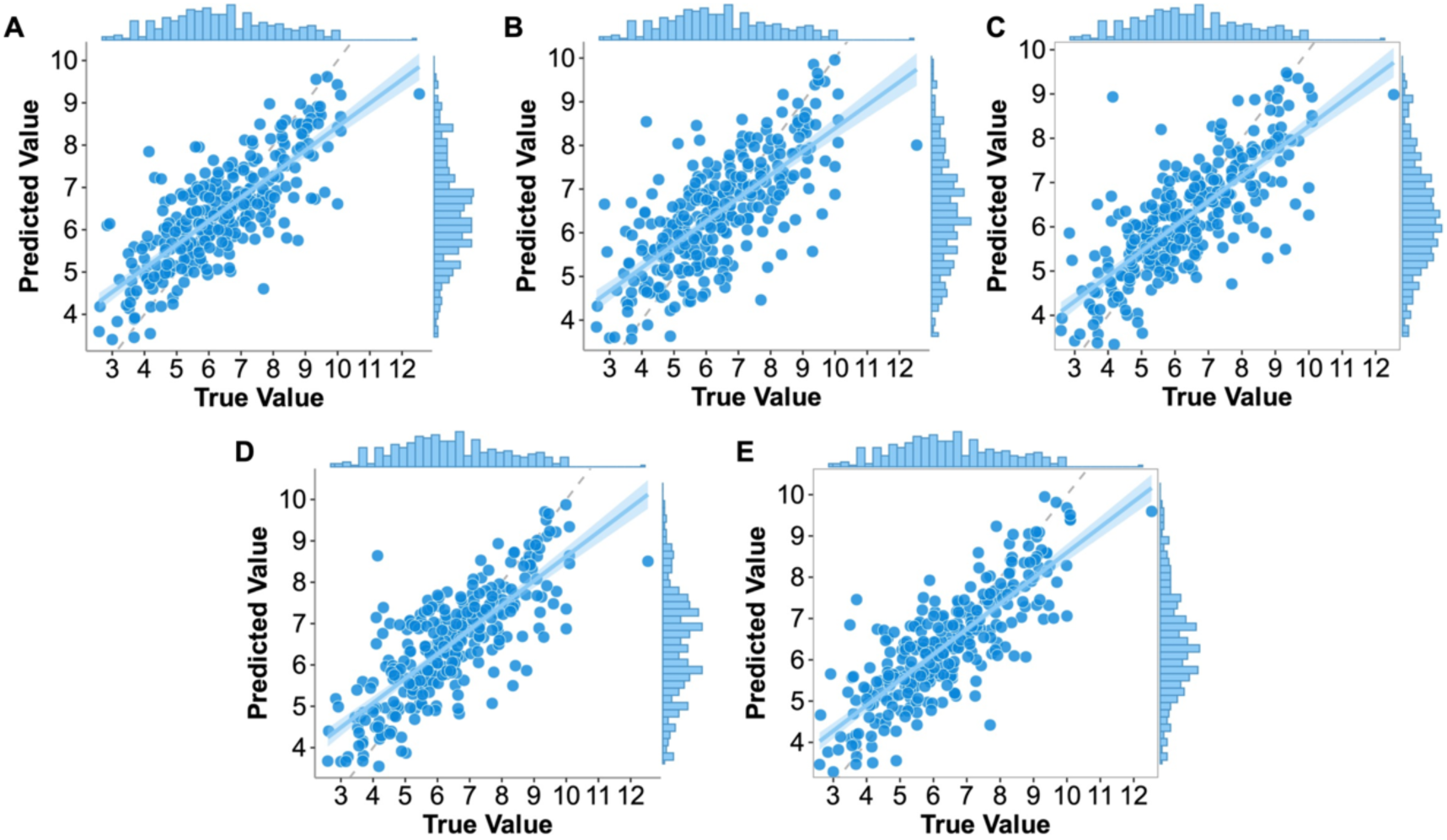
Comparison of predicted and true binding affinities across different models. **A)** XGBoost model; B) DeepDTA; C) TEFDTA D) AttentionDTA; E) PepInter.

### Performance Evaluation of Protein-Peptide Affinity Prediction

Protein-peptide binding affinity is a key determinant of biological activity, interaction specificity and downstream functional outcomes. Accurate prediction of protein-peptide affinity holds significant scientific importance and practical value. In the section, we conducted a systematic assessment using protein-peptide affinity dataset. Similar to the binary interaction prediction setting, all baseline methods (DeepDTA [42], TEFDTA [43], AttentionDTA [44] and XGBoost) were locally installed, trained and tested on the same datasets use five-fold cross-validation to ensure fairness. The comparison results are shown in Table 2. It can be seen that the PepInter achieves the lowest prediction errors among all compared methods, with an RMSE of 1.015 and an MAE of 0.768, indicating a substantial reduction in prediction error. In addition, PepInter attains the highest R² value of 0.654, demonstrating its superior ability to capture the variance in binding affinity data. We also present the scatter plots of different comparison models in Figure 6, the predicted values generally follow the diagonal trend, indicating reasonable predictive performance. However, the predictions generated by PepInter exhibit a noticeably tighter clustering around the diagonal line, accompanied by reduced dispersion compared with the other methods. This observation is consistent with the quantitative improvements reported in Table 2. Moreover, the marginal histograms in each panel reveal that PepInter produces prediction distributions that more closely resemble the distribution of true values, suggesting better calibration and reduced systematic bias. Both the quantitative metrics and qualitative visualizations confirm that PepInter provides more accurate and reliable affinity predictions, highlighting its effectiveness in modeling peptide-protein interactions.

**Table 2.** Performance of PepInter against three methods using five-fold cross-validation on affinity dataset.

| Models | RMSE↓ | MAE↓ | R<sup>2</sup>↑ |
| --- | --- | --- | --- |
| DeepDTA | 1.294±0.059 | 0.983±0.068 | 0.437±0.051 |
| TEFDTA | 1.219±0.059 | 0.907±0.041 | 0.469±0.047 |
| AttentionDTA | 1.209±0.071 | 0.897±0.077 | 0.508±0.058 |
| XGBoost | <u>1.144±0.027</u> | <u>0.871±0.027</u> | <u>0.561±0.021</u> |
| **PepInter** | **1.015±0.028** | **0.768±0.026** | **0.654±0.022** |

### Case study

#### Identifying the PTH type 1 receptor as the target of PTH and PTHrP analogs

Parathyroid hormone receptor type 1 (PTH1R) is a member of the G protein-coupled receptor family and is widely expressed *in vivo*, where it participates in multiple biological processes, including systemic metabolism and tumorigenesis [45]. Parathyroid hormone (PTH) and parathyroid hormone-related peptide (PTHrP) are key regulators of calcium and phosphate metabolism in the body. They also play important roles in the treatment of osteoporosis by increasing bone mass, promoting bone formation and reducing the incidence of high-risk fractures. There have 5 PTH and PTHrP analogs are known to bind to PTH1R [18]. The same procedure described above is applied to remove protein and peptide sequences with sequence similarity greater than 40%. Based on this filtering strategy, the remaining 2,922 proteins were paired with the PTH and PTHrP analogs to construct an independent test set, comprising a total of 14,615 protein-peptide pairs. Table 3 showed the predicted probability between PTH analogs and PTH1R, PepInter was able to predict identify interacting pairs with high AUC value of 0.9662. The PTH1R was consistently placed within the top 5% of candidate proteins for all PTH analogs peptides. These results indicate the strong predictive capability of our model that accurate prediction results can provide valuable risk assessment guidance for pharmacologists in clinical decision-making.

**Table 3.** Predicted interaction probabilities between PTH analogs and PTH1R.

| PDB | Peptide Sequence | Probability |
| --- | --- | --- |
| 1BWX | SVSEIQLMHNLGKHLNSMERVEWLRKKLQDVHNF | 0.996 |
| 6FJ3 | XVAEIQLMHQRAKWLNSLERVEWLRKKLQDVHNYX | 1.000 |
| 6NBF | AVAEIQLMHQRAKWIQDARRRAFLHKLIAEIHTAEI | 1.000 |
| 7VVJ | AVSEHQLLHDKGKSIQDLRRRFFLHHLIAEIHTAEI | 0.887 |
| 8FLS | AVSEHQLLHDKGKSIQDLRRRELLEKLLXKLHTA | 0.996 |

#### Identifying the GLP-1 receptor as the target of semaglutide

The glucagon-like peptide-1 receptor (GLP-1R) is one of the most effective therapeutic targets for the treatment of type 2 diabetes mellitus [46]. It can effectively activate multiple downstream signaling pathways, including PKA, PI3K and MAPK, and is involved in important physiological processes such as insulin secretion and β-cell proliferation [47]. Therefore, GLP-1R, as the functional target through which GLP-1 exerts its effects, has potential significance for guiding the study of multiple diseases. We detected if PepInter was able to identify the interaction of Semaglutide and its analogs with GLP-1R correctly. Following with previous work [33], there exists seven Semaglutide-analogous peptides bind to GLP-1R. To avoid data leakage during the training process, we remove those related peptide drugs from training set which have similar sequence (similarity > 40%) with Semaglutide and remove the protein similar to GLP-1R (similarity > 40%). Afterwards, we combined the rest 2940 proteins with Semaglutide-analogous peptides to build an independent test set that contain 20580 candidate pairs. We then fine tuning our model and predict the results. Table 4 showed the predicted probability of Semaglutide-analogs peptides and GLP-1R pairs, PepInter was able to predict identify interacting pairs with high AUC value of 0.982. The GLP-1R was consistently placed within the top 5% of candidate proteins for all Semaglutide-analogous peptides. We further visualized the prediction results using UMAP in Figure 8, which demonstrates that our model can effectively predict the interactions between Semaglutide-analogous peptides and GLP-1R. These results indicate the strong predictive capability of our model.

**Fig. 8.**
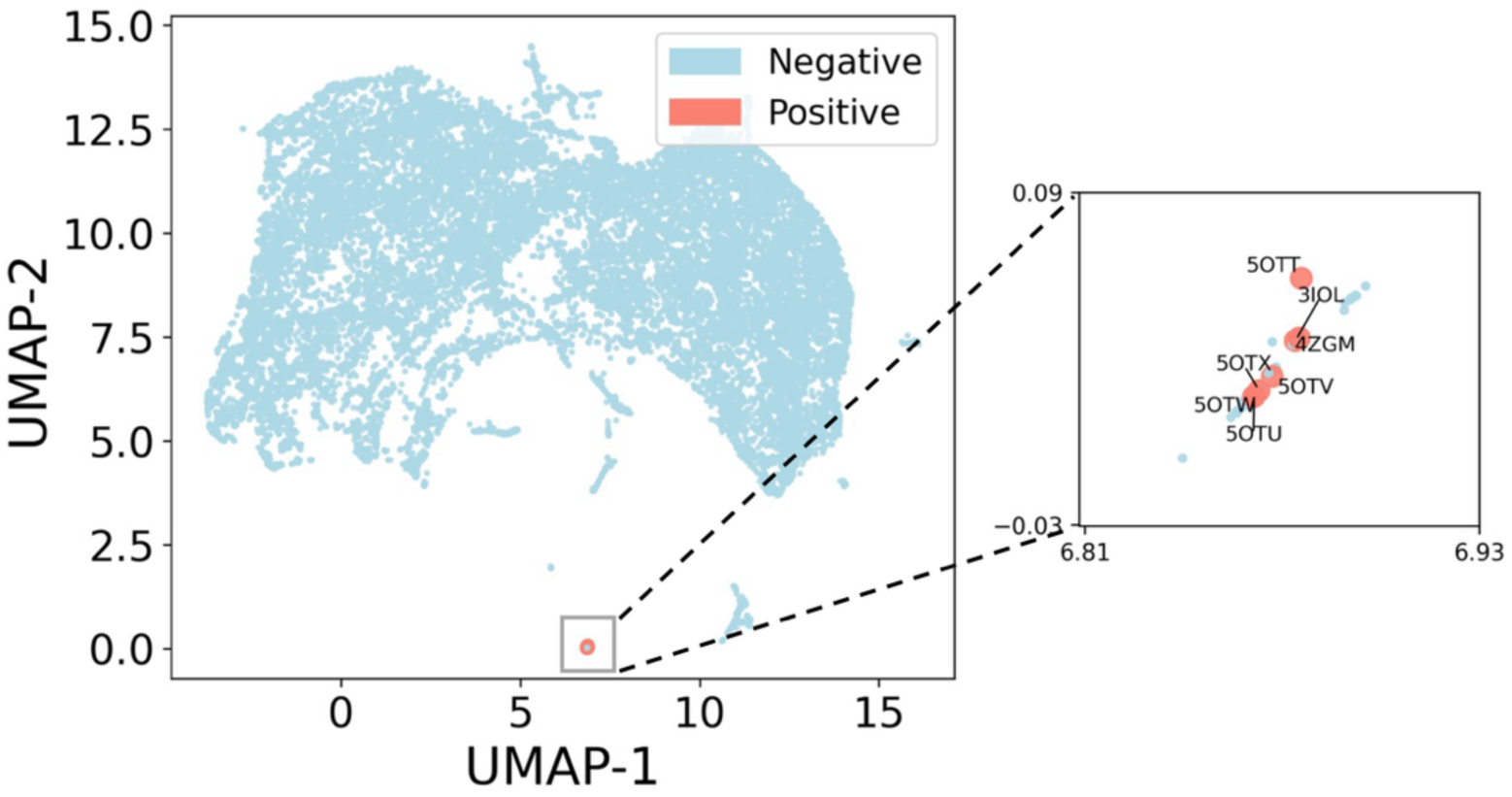
UMAP visualization of predicted interactions between Semaglutide-analogous peptides and GLP-1R. (Light blue and salmon points indicate negative and positive samples, respectively.)

**Table 4.** Predicted interaction probabilities between Semaglutide analogs and GLP-1R.

| PDB | Peptide Sequence | Probability |
| --- | --- | --- |
| 3IOL | HAEGTFTSDVSSYLEGQAAKEFIAWLVKGRG | 0.988 |
| 4ZGM | HAEGTFTSDVSSYLEGQAAKEFIAWLVRGRG | 0.984 |
| 5OTT | HXEGXFTSDLSKQMEEEAVRLFIEWLKNGGPSSGAPPPS | 0.992 |
| 5OTU | HXEGXFTSDVSSYLEGQAAKEFIAWLVKGRG | 0.996 |
| 5OTV | HCEGXFTSDVSSYLEGQAAKEFIAWLVKGRG | 0.996 |
| 5OTW | HXEGCFTSDVSSYLEGQAAKEFIAWLVKGRG | 1.000 |
| 5OTX | HCEGCFTSDVSSYLEGQAAKEFIAWLVKGRG | 1.000 |

## Discussion

PpIs play crucial roles in immune regulation, signal transduction and peptide drug design, efficient and accurate prediction methods are of great importance for cutting-edge biomedical research. However, compared with traditional protein-protein interactions, PpIs exhibit higher structural complexity and are limited by the scale of available experimental data, posing significant challenges for computational prediction approaches. In recent years, extensive efforts have been devoted to PpI prediction. These methods adopt various frameworks and primarily focus on extracting informative representations of proteins and peptides, such as sequence features and structural features. Moreover, Transformer-based and BERT-based prediction frameworks have rapidly advanced, with ESM2 and BERT-related modules demonstrating strong capability in learning structured features directly from protein sequences.

Despite the promising performance achieved by existing methods across multiple datasets, there remains substantial room for improvement before they can fully meet real-world application demands. To address this gap, the large-scale protein-peptide pre-training framework proposed in this study achieves significant performance gains across multiple downstream tasks, validating the effectiveness of pseudo protein-peptide data derived from structural sources and the cross-chain representation learning strategy. The diverse peptide fragments generated through rosetta energy constraints enable the model to learn more comprehensive interfacial energy features, thereby enhancing its ability to characterize binding determinants. The model’s capacity to capture cross-chain contextual dependencies during pre-training further allows more accurate identification of potential binding regions and key peptide motifs, which also explains why the model maintains structural sensitivity even when provided with single-chain sequence input. Moreover, the robust performance observed under imbalanced real-world scenarios indicates that the model not only learns sequence co-occurrence patterns but also infers functional residues and interfacial energy signals.

Compared with traditional methods that rely solely on sequence similarity or local convolutional strategies, our approach retains superior performance when generalizing to unseen proteins or peptides, demonstrating stronger cross-domain transferability-an essential property for peptide drug virtual screening and lead compound optimization. Nevertheless, this study still has some limitations: (i) the current model relies heavily on pseudo energy labels in the pre-training data, and its generalizability to real experimental affinity requires further assessment; (ii) although the model can identify important residues, the underlying binding mechanisms are not yet clearly interpretable; (iii) the sliding-window strategy employed during data construction may overlook long-range binding modes arising from conformational changes; and (iv) explicit structural modeling was not incorporated, suggesting that future work may benefit from integrating AlphaFold-based structure prediction and 3D graph neural networks to enhance geometric awareness.

In the future, we will focus on this work in the following directions: (i) integrating 3D structural constraints and dynamic simulation data to improve the modeling of binding mechanisms; (ii) introducing automated hyperparameter optimization and continual pre-training strategies to enhance the model’s adaptability to new structural domains; and (iii) applying the proposed framework to peptide drug virtual screening and binding affinity optimization, combined with wet-lab experiments to validate its biological efficacy. Overall, this study demonstrates the feasibility of leveraging large-scale pseudo-structural data and cross-chain language modeling to address data scarcity in protein-peptide tasks, providing new insights and technical pathways for computational peptide drug design.

## Conclusion

In this paper, we report a novel sequence-based DL framework, called PepInter, for predicting PpIs. We construct pesudo protein-peptide pairs for learning the contextual dependencies during pre-training stage. and subsequently fine-tune the pre-trained model on downstream PpI prediction tasks using experimentally validated datasets. The comparison with the state-of-the-art methods demonstrates that our method has significantly improved the performance. These outstanding results show that our proposed model PepInter can better capture the latent feature between protein and peptide. Future work will focus on incorporating additional structural information and experimental data to further enhance the predictive performance of PepInter in practical applications. We hope that this approach can accelerate protein-peptide based drug screening and improve the success rate of peptide drug discovery.

## Data availability

Codes will be update in Github.

## References

1. Petsalaki, E.; Russell, R. B. Peptide-Mediated Interactions in Biological Systems: New Discoveries and Applications. Curr. Opin. Biotechnol. 2008, 19, 344–350.

2. Huang, Y.; He, L.; Li, G.; Zhai, N.; Jiang, H.; Chen, Y. Role of helicity of α-helical antimicrobial peptides to improve specificity. *Prot*. Cell, 2014, 5, 631–642.

3. Kobayashi, M.; Onozawa, N.; Matsuda, K.; Wakimoto, T. Chemoenzymatic tandem cyclization for the facile synthesis of bicyclic peptides. Comm. Chem. 2024, 7, 67.

4. Hong, R.; Xie, A.; Jiang, C.; Guo, Y.; Zhang, Y.; Chen, J.; Shen, X.; Li, M.; Yue, X. A review of the biological activities of lactoferrin: Mechanisms and potential applications. Food Funct. 2024, 15, 8182–8199.

5. Shi, C.L.; Von Wangenheim, D.; Herrmann, U.; Wildhagen, M.; Kulik, I.; Kopf, A.; Ishida, T.; Olsson, V.; Anker, M.K.; Albert, M.; Butenko, M.A. The dynamics of root cap sloughing in Arabidopsis is regulated by peptide signalling. Nat. Plants. 2018, 4, 596–604.

6. Dyson, H.J.; Wright, P.E. Intrinsically unstructured proteins and their functions. Nat. Rev. Mol. Cell. Biol. 2005, 6, 197–208.

7. Cunningham, J.M.; Koytiger, G.; Sorger, P.K.; AlQuraishi, M. Biophysical prediction of protein–peptide interactions and signaling networks using machine learning. Nat. Met. 2020, 17, 175–183.

8. Shanker, S.; Sanner, M.F. Predicting protein-peptide interactions: Benchmarking deep learning techniques and a comparison with focused docking. J. Chem. Inf. Model. 2023, 63, 10, 3158–3170.

9. Kozlovskii, I.; Popov, P. Protein-peptide binding site detection using 3D convolutional neural networks. J. Chem. Inf. Model. 2021, 61, 3814–3823.

10. Shafiee, S.; Fathi, A.; Taherzadeh, G. SPPPred: sequence-based protein-peptide binding residue prediction using genetic programming and ensemble learning. IEEE ACM. Trans. Comput. Bi. 2022, 20(3), pp.2029–2040.

11. Zhou, P.; Jin, B.; Li, H.; Huang, S.Y. HPEPDOCK: a web server for blind peptide–protein docking based on a hierarchical algorithm. Nucleic Acids Res. 2018, 46, W443–W450.

12. London, N.; Raveh, B.; Cohen, E.; Fathi, G.; Schueler-Furman, O. Rosetta FlexPepDock web server-high resolution modeling of peptide-protein interactions. Nucleic Acids Res. 2011, 39, W249–W253.

13. de Vries, S.J.; Rey, J.; Schindler, C.E.; Zacharias, M.; Tuffery, P. The pepATTRACT web server for blind, large-scale peptide-protein docking. Nucleic Acids Res. 2017, 45, W361–W364.

14. Taherzadeh, G.; Zhou, Y.; Liew, A.W.C.; Yang, Y. Structure-based prediction of protein–peptide binding regions using Random Forest. Bioinform. 2018, 34, 477–484.

15. Das, A.A.; Sharma, O.P.; Kumar, M.S.; Krishna, R.; Mathur, P.P. PepBind: a comprehensive database and computational tool for analysis of protein-peptide interactions. Genom. Proteom. Bioinform. 2013, 11(4), pp.241–246.

16. Johansson-Åkhe, I.; Mirabello, C.; Wallner, B. Predicting protein-peptide interaction sites using distant protein complexes as structural templates. Sci. rep. 2019, 9, 4267.

17. Chen, S.; Yan, K.; Li, X.; Liu, B. Protein language pragmatic analysis and progressive transfer learning for profiling peptide–protein interactions. IEEE Trans. Neural Networks Learn. Syst. 2025, 8, 15385–15399.

18. Zhou, R.; Zhao, H.; Zhong, J.; Duan, G. November. PepGPL: A Multi-Task Framework for Identifying Peptide-Protein Interactions and Corresponding Binding Residues. International Conference on Bioinformatics, Computational Biology and Health Informatics. 2024, 22, 1–9.

19. Sun, X.; Wu, Z.; Su, J.; Li, C. A deep attention model for wide-genome protein-peptide binding affinity prediction at a sequence level. Int. J. Biol. Macromol. 2024, 276, 133811.

20. Beal, D.J. ESM 2.0: State of the art and future potential of experience sampling methods in organizational research. Annu. Rev. Organ. Psychol. Organ. Behav. 2015, 2, 383–407.

21. Koroteev, M.V.; BERT: a review of applications in natural language processing and understanding. arXiv preprint arXiv:2103.11943, 2021.

22. Elnaggar, A.; Heinzinger, M.; Dallago, C.; Rehawi, G.; Wang, Y.; Jones, L.; Gibbs, T.; Feher, T.; Angerer, C.; Steinegger, M.; Bhowmik, D. ProtTrans: Toward understanding the language of life through self-supervised learning. IEEE Trans. Pattern Anal. Mach. Intell. 2021, 44, 7112–7127.

23. Jin, X.; Chen, Z.; Yu, D.; Jiang, Q.; Chen, Z.; Yan, B.; Qin, J.; Liu, Y.; Wang, J. TPepPro: a deep learning model for predicting peptide-protein interactions. Bioinf. 2025, 41, btae708.

24. London, N.; Raveh, B.; Schueler-Furman, O. Druggable protein–protein interactions–from hot spots to hot segments. Curr. Opin. Biotechnol. 2013, 17, 952–959.

25. Sedan, Y.; Marcu, O.; Lyskov, S.; Schueler-Furman, O. Peptiderive server: derive peptide inhibitors from protein–protein interactions. Nucleic Acids Res. 2016, 4, W536–W541.

26. Dapkūnas, J.; Timinskas, A.; Olechnovič, K.; Tomkuvienė, M.; Venclovas, Č. PPI3D: a web server for searching, analyzing and modeling protein-protein, protein-peptide and protein-nucleic acid interactions. Nucleic Acids Res. 2024, 52, W264–W271.

27. Steinegger, M.; Söding, J. MMseqs2 enables sensitive protein sequence searching for the analysis of massive data sets. Nat. Biotechnol. 2017, 35, 1026–1028.

28. Omidi, A.; Møller, M.H.; Malhis, N.; Bui, J.M.; Gsponer, J. AlphaFold-Multimer accurately captures interactions and dynamics of intrinsically disordered protein regions. Proc. Natl. Acad. Sci. 2024, 121, e2406407121.

29. Wang, R.; Fang, X.; Lu, Y.; Wang, S.; The PDBbind database: Collection of binding affinities for protein− ligand complexes with known three-dimensional structures. J. Med. Chem. 2004, 47, 2977–2980.

30. Jankauskaitė, J.; Jiménez-García, B.; Dapkūnas, J.; Fernández-Recio, J.; Moal, I.H. SKEMPI 2.0: an updated benchmark of changes in protein-protein binding energy, kinetics and thermodynamics upon mutation. Bioinfor. 2019, 35, 462–469.

31. Dunbar, J.; Krawczyk, K.; Leem, J.; Baker, T.; Fuchs, A.; Georges, G.; Shi, J.; Deane, C.M. SAbDab: the structural antibody database. Nucleic Acids Res. 2014, 42, D1140–D1146.

32. He, Y.; Tian, Z.; Zheng, J.; Wang, H.; Han, L.; Han, W. Prediction of Umami Peptides Based on a Large Language Model of Proteins. J. Chem. Inf. Model. 2025, 65, 3955–3962.

33. Lei, Y.; Li, S.; Liu, Z.; Wan, F.; Tian, T.; Li, S.; Zhao, D.; Zeng, J. A Deep-Learning Framework for Multi-Level Peptide-Protein Interaction Prediction. Nat. Commun. 2021, 12.

34. ESM Team. ESM Cambrian: Revealing the mysteries of proteins with unsupervised learning. EvolutionaryScale Website, 2024, https://evolutionaryscale.ai/blog/esm-cambrian.

35. Lin, T.Y.; Goyal, P.; Girshick, R.; He, K.; Dollár, P. Focal loss for dense object detection. IEEE Int. Conf. Comput. Vis. 2017, 2980–2988.

36. Makansi, O.; Ilg, E.; Cicek, O.; Brox, T. Overcoming limitations of mixture density networks: A sampling and fitting framework for multimodal future prediction. IEEE Comput. Soc. Conf. Comput. Vis. Pattern Recognit. 2019, 7144–7153.

37. Loshchilov, I.; Hutter, F. Decoupled weight decay regularization. arXiv preprint arXiv:1711.05101. 2017.

38. Chen, T. XGBoost: A Scalable Tree Boosting System. Cornell University, 2016.

39. Zhan, X.; Liu, T.; Yu, C.; Huang, Y.A.; You, Z.; Siu, S.W. MAARDTI: a multi-perspective attention aggregation model for the prediction of drug–target interactions. Digi. Discov. 2025, 4, 2994–3007.

40. Zhao, Q.; Zhao, H.; Zheng, K.; Wang, J. HyperAttentionDTI: improving drug–protein interaction prediction by sequence-based deep learning with attention mechanism. Bioinfor. 2022, 38, 655–662.

41. Deng, M.; Wang, J.; Zhao, Y.; Zhao, Y.; Cao, H.; Wang, Z. Predicting drug and target interaction with dilated reparameterize convolution. Sci. Rep. 2025, 15, 2579.

42. Öztürk, H.; Özgür, A.; Ozkirimli, E. DeepDTA: deep drug-target binding affinity prediction. Bioinfor. 2018, 34, i821–i829.

43. Li, Z.; Ren, P.; Yang, H.; Zheng, J.; Bai, F. TEFDTA: a transformer encoder and fingerprint representation combined prediction method for bonded and non-bonded drug-target affinities. Bioinfor. 2024, 40, btad778.

44. Zhao, Q.; Duan, G.; Yang, M.; Cheng, Z.; Li, Y.; Wang, J. AttentionDTA: Drug–target binding affinity prediction by sequence-based deep learning with attention mechanism. IEEE/ACM Trans. Comput. Biol. Bioinform. 2022, 20, 852–863.

45. Martin, T.J.; Sims, N.A.; Seeman, E. Physiological and pharmacological roles of PTH and PTHrP in bone using their shared receptor, PTH1R. Endoc. Rev. 2021, 42, 383–406.

46. Blad, C.C.; Tang, C.; Offermanns, S. G protein-coupled receptors for energy metabolites as new therapeutic targets. Nat. Rev. Drug Discov. 2012, 11, 603–619.

47. Koole, C.; Savage, E.E.; Christopoulos, A.; Miller, L.J.; Sexton, P.M.; Wootten, D. Minireview: signal bias, allosterism, and polymorphic variation at the GLP-1R: implications for drug discovery. Mole. Endocrinol. 2013, 27, 1234–1244.

48. Engelhardt, P.M.; Florez-Rueda, S.; Drexelius, M.; Neudörfl, J.M.; Lauster, D.; Hackenberger, C.P.; Kühne, R.; Neundorf, I.; Schmalz, H.G. Synthetic α-Helical Peptides as Potential Inhibitors of the ACE2 SARS-CoV-2 Interaction. ChemBioChem, 2022, 23, e202200372.

49. Li, Y.; Pan, D.; Zhang, W.; Xie, X.; Dang, Y.; Gao, X. Identification and Molecular Mechanism of Novel ACE Inhibitory Peptides from Broccoli Protein. Food Biosci. 2024, 61, 104678.

