## Supplementary material for "Protein-peptide Interaction Representation Learning with Pretrained Language Models": SI

---

### The parameter of the Rosetta Peptiderive protocol

**Table S1.** Hyperparameter setting in the Rosetta Peptiderive protocol.

| Option name | Type | Setting |
|---|---|---|
| pep_lengths | List of numbers | **5-50** |
| skip_zero_isc | True/False | True |
| dump_peptide_pose | True/False | True |
| dump_cyclic_poses | True/False | False |
| dump_prepared_pose | True/False | True |
| dump_report_file | True/False | True |
| restrict_receptors_to_chains | List of characters | - |
| restrict_partners_to_chains | List of characters | - |
| do_minimize | True/False | True |
| optimize_cyclic_threshold | Real number | 0.35 |
| report_format | String | Markdown |

### Baseline Methods

In order to better evaluate our proposed method, we make a comparison with several following competitive methods for predicting PpI and affinity. To ensure the fairness of comparison, all the baseline models are trained and evaluated on the same datasets and most hyper-parameters are set based on the original works.

The comparison models for predicting PpI are list as follows:

- **XGBoost**: XGBoost was an efficient and powerful ensemble decision tree model used to learn patterns from features and make predictions. For the XGBoost baseline, protein and peptide sequences were first encoded using the pretrained ESM2 model, and the resulting embeddings were concatenated to form joint representations, which were then used as input to an XGBoost model with hyperparameters optimized using Optuna [] for binary classification.
- **MAARDTI**: MAARDTI is used to predict drug-target interaction. It utilizes a multi-layer CNN block to learn low-dimensional representation features. The multi-perspective attention aggregating module and bi-contextual refocusing module are adopted to learn the interaction relationship between proteins and drugs, the combined features are finally fed into interaction prediction block. In this study, drug embedding is replaced with a peptide embedding while keeping the rest of the framework unchanged.
- **Rep-ConvDTI**: Rep-ConvDTI is a deep learning framework for drug-target interaction prediction that models large-scale sequence patterns using large-kernel convolutions,

while capturing fine-grained information through reparameterization and a gated attention mechanism to better characterize drug–target interactions. In this study, we modify the target encoding to peptide encoding, without altering the overall framework of Rep-ConvDTI.

- **HyperAttentionDTI**: HyperAttentionDTI utilizes the attention mechanism to capture attention vector of each amino acid and atom. This model use CNN block to extract protein and drug features, and mixed original feature and HyperAttention matrix. The modified drug-protein pair feature vector is fed into FCN for final predictions. In this model, we replace the drug encoding with a peptide encoding, while keeping all other hyperparameter settings consistent with the original work.
- **PIPR**: PIPR is an end-to-end sequence-based protein-protein interaction prediction model that uses a Siamese residual RCNN to automatically learn both local and contextual features from protein sequences, achieving strong performance without manual feature engineering.

The comparison models for predicting affinity are list as follows:

- **XGBoost:** For the XGBoost baseline, protein and peptide sequences were first encoded using the pretrained ESM2 model, and the resulting embeddings were concatenated to form joint representations. For affinity prediction, the model was adapted to a regression setting by modifying only the output layer, while keeping all other components unchanged, with hyperparameters optimized using Optuna.
- **DeepDTA**: DeepDTA utilizes two parallel 3-layer CNNs and fully connected layers for predicting DTI. It originally extracts drug and protein feature maps by CNN block and then drug and protein feature maps are concatenated. Finally, the model obtained prediction result by fed drug-target pairs into fully connected layers.
- **TEFDTA**: TEFDTA is an attention-based framework for drug-target binding affinity prediction that integrates a Transformer encoder with molecular fingerprints to model both non-covalent and covalent interactions. By employing distinct representations for proteins and drugs and leveraging fine-tuning from large non-covalent datasets to smaller covalent datasets, TEFDTA effectively improves affinity prediction accuracy across different interaction types.
- **AttentionDTA**: AttentionDTA is an end-to-end deep learning framework for drug-target binding affinity prediction that integrates attention mechanisms with one-dimensional convolutional neural networks to jointly model drug and protein sequences. By explicitly capturing the mutual importance of subsequences between drugs and targets, AttentionDTA learns more informative representations and improves affinity prediction performance.

For the fairness of the experiment, all hyperparameters are kept consistent with those used in the original work.

#### Comparative performance of PepInter against baseline methods under different clustering settings

**Table S2.** Performance of PepInter against fivemethods under different clustering settings.

| Model_Name | Threshold | Type | ACC | REC | F1 | AUC |
|---|---|---|---|---|---|---|
| PepInter | 0.3 | pair | 0.941 | 0.939 | 0.942 | 0.980 |
| | 0.5 | pair | 0.946 | 0.935 | 0.944 | 0.982 |
| | 0.8 | pair | 0.941 | 0.942 | 0.940 | 0.982 |
| | 1 | pair | 0.944 | 0.939 | 0.942 | 0.983 |
| | 0.3 | protein | 0.892 | 0.868 | 0.878 | 0.937 |
| | 0.5 | protein | 0.905 | 0.837 | 0.881 | 0.965 |
| | 0.8 | protein | 0.919 | 0.870 | 0.908 | 0.968 |
| | 1 | protein | 0.921 | 0.871 | 0.910 | 0.969 |
| | 0.3 | peptide | 0.919 | 0.892 | 0.914 | 0.973 |
| | 0.5 | peptide | 0.928 | 0.904 | 0.925 | 0.975 |
| | 0.8 | peptide | 0.931 | 0.913 | 0.927 | 0.978 |
| | 1 | peptide | 0.931 | 0.913 | 0.927 | 0.978 |
| XGBoost | 0.3 | pair | 0.903 | 0.890 | 0.901 | 0.963 |
| | 0.5 | pair | 0.916 | 0.905 | 0.912 | 0.968 |
| | 0.8 | pair | 0.920 | 0.914 | 0.917 | 0.975 |
| | 1 | pair | 0.924 | 0.920 | 0.919 | 0.978 |
| | 0.3 | protein | 0.848 | 0.751 | 0.811 | 0.923 |
| | 0.5 | protein | 0.863 | 0.794 | 0.830 | 0.935 |
| | 0.8 | protein | 0.875 | 0.837 | 0.860 | 0.942 |
| | 1 | protein | 0.882 | 0.841 | 0.862 | 0.945 |
| | 0.3 | peptide | 0.899 | 0.871 | 0.893 | 0.959 |
| | 0.5 | peptide | 0.901 | 0.879 | 0.896 | 0.963 |
| | 0.8 | peptide | 0.908 | 0.894 | 0.904 | 0.965 |
| | 1 | peptide | 0.910 | 0.901 | 0.907 | 0.964 |
| PIPR | 0.3 | pair | 0.839 | 0.831 | 0.837 | 0.907 |
| | 0.5 | pair | 0.847 | 0.839 | 0.842 | 0.910 |
| | 0.8 | pair | 0.841 | 0.828 | 0.836 | 0.911 |
| | 1 | pair | 0.845 | 0.831 | 0.829 | 0.914 |
| | 0.3 | protein | 0.723 | 0.676 | 0.679 | 0.806 |
| | 0.5 | protein | 0.728 | 0.614 | 0.654 | 0.783 |

| | | | | | | |
|---|---|---|---|---|---|---|
| | 0.8 | protein | 0.760 | 0.740 | 0.739 | 0.837 |
| | 1 | protein | 0.767 | 0.752 | 0.743 | 0.841 |
| | 0.3 | peptide | 0.808 | 0.809 | 0.803 | 0.880 |
| | 0.5 | peptide | 0.825 | 0.821 | 0.820 | 0.891 |
| | 0.8 | peptide | 0.717 | 0.827 | 0.729 | 0.819 |
| | 1 | peptide | 0.802 | 0.830 | 0.731 | 0.820 |
| Rep-ConvDTI | 0.3 | pair | 0.865 | 0.861 | 0.863 | 0.921 |
| | 0.5 | pair | 0.859 | 0.871 | 0.857 | 0.914 |
| | 0.8 | pair | 0.866 | 0.882 | 0.865 | 0.930 |
| | 1 | pair | 0.872 | 0.892 | 0.869 | 0.931 |
| | 0.3 | protein | 0.767 | 0.642 | 0.705 | 0.796 |
| | 0.5 | protein | 0.794 | 0.717 | 0.745 | 0.825 |
| | 0.8 | protein | 0.803 | 0.726 | 0.772 | 0.861 |
| | 1 | protein | 0.811 | 0.735 | 0.793 | 0.892 |
| | 0.3 | peptide | 0.834 | 0.800 | 0.824 | 0.880 |
| | 0.5 | peptide | 0.816 | 0.768 | 0.803 | 0.886 |
| | 0.8 | peptide | 0.840 | 0.823 | 0.832 | 0.905 |
| | 1 | peptide | 0.862 | 0.847 | 0.520 | 0.918 |
| HyperAttentionDTI | 0.3 | pair | 0.890 | 0.876 | 0.888 | 0.946 |
| | 0.5 | pair | 0.894 | 0.876 | 0.889 | 0.944 |
| | 0.8 | pair | 0.897 | 0.859 | 0.891 | 0.956 |
| | 1 | pair | 0.908 | 0.871 | 0.899 | 0.958 |
| | 0.3 | protein | 0.800 | 0.662 | 0.742 | 0.862 |
| | 0.5 | protein | 0.799 | 0.657 | 0.733 | 0.863 |
| | 0.8 | protein | 0.831 | 0.705 | 0.793 | 0.898 |
| | 1 | protein | 0.843 | 0.716 | 0.805 | 0.912 |
| | 0.3 | peptide | 0.852 | 0.788 | 0.837 | 0.913 |
| | 0.5 | peptide | 0.859 | 0.819 | 0.850 | 0.929 |
| | 0.8 | peptide | 0.860 | 0.852 | 0.854 | 0.936 |
| | 1 | peptide | 0.882 | 0.865 | 0.863 | 0.940 |
| MAARDTI | 0.3 | pair | 0.887 | 0.858 | 0.883 | 0.945 |
| | 0.5 | pair | 0.889 | 0.860 | 0.883 | 0.936 |
| | 0.8 | pair | 0.887 | 0.862 | 0.883 | 0.941 |
| | 1 | pair | 0.899 | 0.874 | 0.892 | 0.944 |
| | 0.3 | protein | 0.791 | 0.662 | 0.745 | 0.845 |
| | 0.5 | protein | 0.790 | 0.606 | 0.748 | 0.860 |
| | 0.8 | protein | 0.833 | 0.749 | 0.813 | 0.900 |
| | 1 | protein | 0.845 | 0.758 | 0.821 | 0.911 |
| | 0.3 | peptide | 0.847 | 0.808 | 0.838 | 0.906 |
| | 0.5 | peptide | 0.860 | 0.844 | 0.855 | 0.920 |

| | | | | | | |
|---|---|---|---|---|---|---|
| | 0.8 | peptide | 0.877 | 0.866 | 0.871 | 0.928 |
| | 1 | peptide | 0.884 | 0.874 | 0.884 | 0.929 |
